# Stochastic regimes can hide the attractors in data, reconstruction algorithms can reveal them

**DOI:** 10.1101/2024.02.17.580797

**Authors:** Babak M. S. Arani, Stephen R. Carpenter, Egbert H. van Nes, Ingrid A. van de Leemput, Chi Xu, Pedro G. Lind, Marten Scheffer

## Abstract

Tipping points and alternative attractors have become an important focus of research and public discussions about the future of climate, ecosystems and societies. However, empirical evidence for the existence of alternative attractors remains scarce. For example, bimodal frequency distributions of state variables may suggest bistability, but can also be due to bimodality in external conditions. Here, we bring a new dimension to the classical arguments on alternative stable states and their resilience showing that the stochastic regime can distort the relationship between the probability distribution of states and the underlying attractors. Simple additive Gaussian white noise produces a one-to-one correspondence between the modes of frequency distributions and alternative stable states. However, for more realistic types of noise, the number and position of modes of the frequency distribution do not necessarily match the equilibria of the underlying deterministic system. We show that data must represent the stochastic regime as thoroughly as possible. When data are adequate then existing methods can be used to determine the nature of the underlying deterministic system and noise simultaneously. This may help resolve the question of whether there are tipping points, but also how realized states of a system are shaped by stochastic forcing vs internal stability properties.

**Open Research Statement:** Data and MATLAB codes for results reported here are available in the Github repository https://github.com/mshoja/Reconst (Babak M. S. Arani 2023) The original data source is cited in the text.

## INTRODUCTION

As nature and humanity face rapid change, ecology, climate science and social sciences are increasingly interested in understanding tipping points, critical thresholds where change becomes self-reinforcing causing it to speed up in sometimes irreversible ways. Focusing on landscapes and ecosystems it is obvious that massive changes happen at multiple space and time scales. A central current challenge is to find out which states of ecosystems and landscapes are likely to persist in the future, at which scales, under what disturbance regimes, with how much and what kinds of intrinsic variation. These concerns prompt the need to identify potentially persistent states from masses of spatial and temporal data even as the earth system seems to be losing stationarity at relevant scales necessary for sustaining civilization. To meet this challenge, we need methods for identifying desirable and potentially persistent states of complex systems in highly variable contexts.

So far it has been difficult to rigorously show that an ecosystem has alternative stable states. Scheffer and Carpenter (Scheffer and Carpenter 2003) suggested six hints from field data and experiments that provides evidence for the existence of alternative ecosystem states, but stressed that none of them is a proof as other explanations remain possible. One of the most commonly used hints from field data is the expectation that the frequency distribution of key variables of systems with multiple stable states is multimodal (Scheffer and Carpenter 2003), where the modes match the underlying equilibria (Livina et al. 2010). This has been applied to different systems, ranging from forests (Hirota et al. 2011, Scheffer et al. 2012, Xu et al. 2016, Moris et al. 2017), human microbiomes (Lahti et al. 2014) to seagrass beds (de Fouw et al. 2016). However, like the other hints from field data, multimodality is not irrefutable evidence for alternative stable states as it can also be due to multimodality in the external conditions (Scheffer and Carpenter 2003).

Here, we discuss other reasons why this analysis may fail to show alternative stable states. The frequency distribution only shows the alternative stable states accurately if two assumptions are met (Livina et al., 2010). The first assumption is that the time span of data is long enough to assume that the data distribution is in equilibrium, the so-called *stationary probability distribution* (Gardiner 1985). The second assumption is that the dynamics can be described by a stochastic differential equation where the stochastic fluctuations can be classified as ‘additive Gaussian white noise’ (Kwasniok and Lohmann 2009, Livina et al. 2010). This is one of the simplest assumptions about the nature of stochastic fluctuations, and is commonly used in applied sciences, either because the true features of the stochasticity are unknown or because this assumption facilitates some mathematical analyses. The key properties of additive Gaussian white noise, as the name implies, include: (1) the intensity of stochastic fluctuations (see Appendix S1) is independent of the state of the system (additive), (2) the stochasticity is drawn from a Gaussian distribution, and (3) there is no temporal autocorrelation in the added stochastic fluctuations (white).

Indeed, for this kind of noise there is a one-to-one correspondence between the positions of alternative stable states (i.e., the state or regime to which a system will asymptotically settle in the absence of perturbations) and the modes of stationary probability distribution (see Appendix S1). However, the assumption of additive Gaussian white noise is highly unrealistic for most, if not all, natural situations (Vasseur and Yodzis 2004).We show that for other more realistic noise regimes the correspondence between the attractors and the modes of probability distributions is partly or completely lost. We suggest that there are other methods that can indicate alternative stable states in those cases.

## METHODS

### The modelling in a nutshell

We use the following one-dimensional Langevin model (Gardiner 1985)

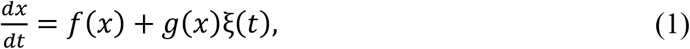

to capture and describe the dynamics reflected in noisy time series data. In (2), the function *f*(*x*) represents the deterministic part of the system as a function of state *x* which can, for instance, be the amount of biomass or population size. ξ(*t*) represents the stochastic fluctuations (the noise). In classical modelling ξ(*t*) is assumed to have a Gaussian distribution and be completely uncorrelated in time (white) which are, indeed, idealizations. The whiteness of noise reflects a ‘fast fluctuating’ environment which allows time scale separation between the deterministic mechanisms and the surrounding environment. This means that the autocorrelation function of a white noise drops immediately. In reality, however, noise often has an autocorrelation structure which will decay based on a finite correlation time. The Gaussian assumption, bearing in mind the Central Limit Theorem (CLT), is based on the idealization that the environmental forces are repetitively drawn from distributions with finite variance. In reality, however, such environmental forces can be drawn from distributions which might have heavy tails (and therefore may not have a finite variance). In such cases a generalized CLT should be used leading to a noise distribution with heavy-tails (unlike the Gaussian distribution which has an exponentially decaying tails). Here, we consider the effects of non-Gaussian and correlated (colored) noises on the number and position of alternative stable states. The function *g*(*x*) describes how the intensity of statistical fluctuations varies with state, so *g*(*x*) can be intuitively understood as a ‘weight’ to the contribution of stochastic fluctuations at each state. If *g*(*x*) does not depend on the state (i.e., it is a constant) then the noise in (3) is said to be ‘additive’, otherwise it is called ‘multiplicative’. Note that for the main purpose of the paper it is sufficient to use a one-dimensional model as we did. Parallel to (4) one can model the evolution of the probability distribution function of states at any time, i.e., *p*(*x, t*) by a partial differential equation (Gardiner 1985). For very large times (*t* → ∞) this distribution might approach a distribution function *p*^*st*^(*x*) which is independent of time and is called ‘stationary distribution’ for which there exists an analytical solution in terms of functions *f*(*x*) and g(*x*) when noise is Gaussian and white (see formula (2) in appendix S1). For more complex noise sources we estimate the stationary distribution numerically via simulation of the system. We refer to the modes (i.e., most visited states) of the stationary distribution as the ‘alternative stable states’ of (5). However, determining the alternative stable states using the classical deterministic methodology by solving for *f*(*x*) = 0 might not coincide with the modes of the stationary distribution when the noise is complex. Below, we clarify what we mean by ‘complex noise’.

### Complex noise in ecological systems

By complex noise, we mean any deviation from additive Gaussian white noise, i.e., either noise is not additive (Figure 1B) or, it follows a non-Gaussian distribution (Figure 1D) or, it has memory (colored noise, which has a temporally autocorrelated structure) (Figure 1F) or any combination of these. Clearly, ecological systems are complex systems, characterized by many variables interacting in complicated ways. A vast number of these variables operate at short time scales with small amplitudes (fast variables, microscopic variables or random forces), relative to a few variables of interest evolving slowly (slow variables or macroscopic variables). One can effectively treat the collective effect of random forces as noise and keep the slow variables only (Haken and Synergetics 2004). Taking forest ecosystems as an example, processes such as temperature fluctuation, wind disturbance and insect grazing evolve much faster than forest biomass accumulation on a macroscopic scale. Ecologists who are interested in the long-term dynamics of forest biomass can treat the overall effect of these fast variables as noise. Viewed this way, noise is nothing but our lack of information about the true state of the system, i.e., ‘system noise’. Note that this should not be confused with ‘measurement noise’ (Scholz et al. 2016).

**Figure 1.**
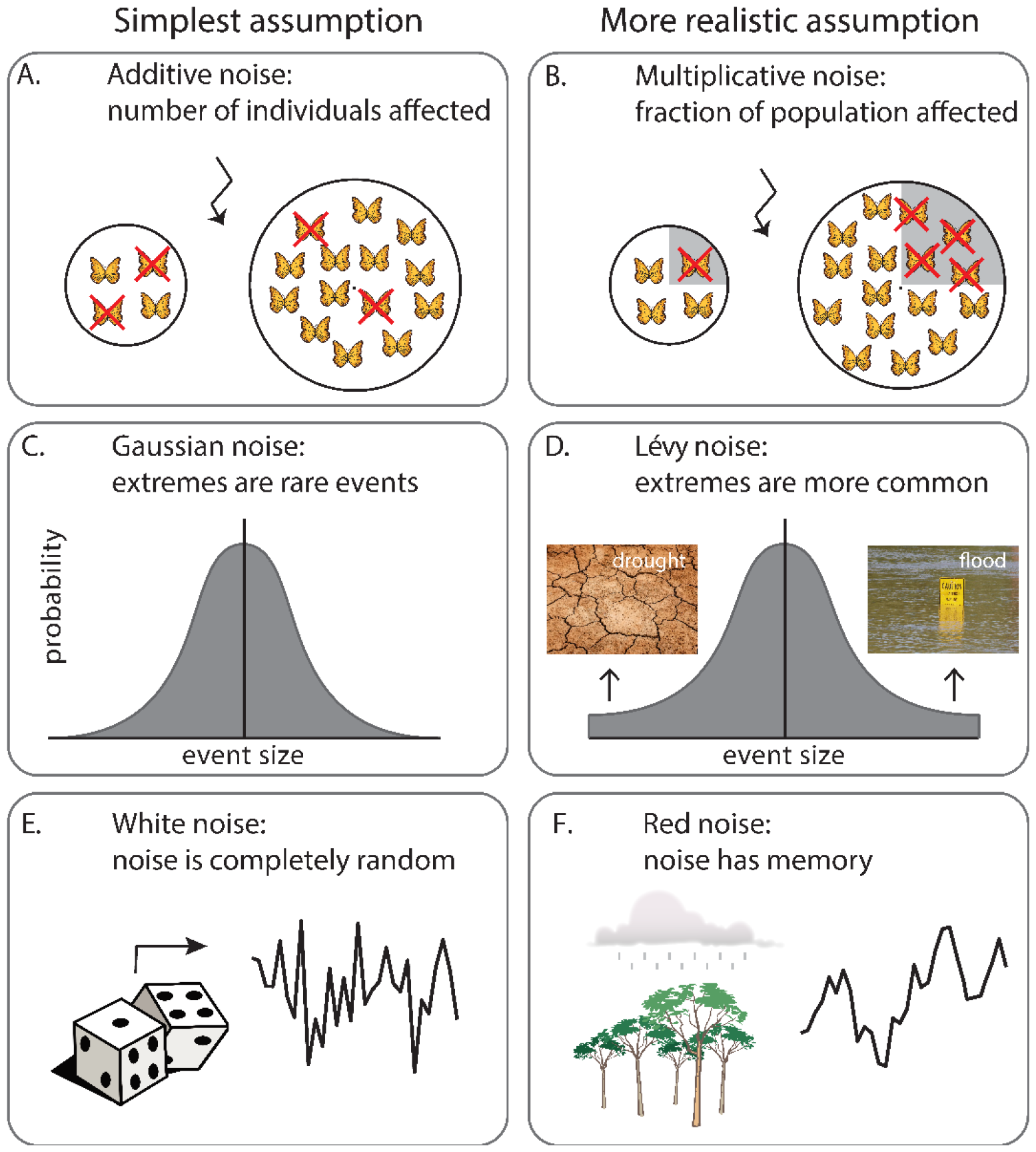
Simple assumptions about the nature of stochasticity (A, C, and E) versus more realistic alternatives (B, D and F). In the context of population dynamics, additive noise can affect a fixed number of individuals (A), whereas multiplicative noise may affect a fraction of population (B). Often, random forces contributing to the noise can, and in essence, belong to a distribution with heavy tails. For instance, flood and drought can be thought of as extremes (D) which, although still rare, can occur more often than normal distribution predicts (C). The overall effect of such random forces (noise) tends to have a stable distribution with asymptotic power-law tails in which the Gaussian distribution is just an extreme member (Generalized central limit Theorem, see appendix S3). To reduce complexity of ecosystems that are high-dimensional systems (characterized by a vast number of variables), ecologists often monitor a few key variables. The resulting low-dimensional system considerably simplifies the analysis of the original high-dimensional one. Such an oversimplification is, however, costly leading to a now low-dimensional system which relies on its past states, i.e., a colored noise source (Hanggi and Jung 1995), (F). The two photographs in Figure 1D are via Pixabay website where the photograph for drought can be found in the link https://pixabay.com/photos/earth-drought-floor-dryness-3355931/ and the one for the flood can be found in the link https://pixabay.com/photos/flood-tennessee-river-damage-17506/

### Inferring stochastic dynamical systems governing the data: system reconstruction

System reconstruction refers to any technique being used to fit a (stochastic) system of dynamical systems to time series data (Siegert et al. 1998, Tabar 2019). In a nutshell, system reconstruction is about disentangling and estimating the deterministic (i.e., *f*(*x*) in (6)) and stochastic (i.e., *g*(*x*) in (7) and properties of the noise source ξ(*t*)) forces in which the interaction between them leads to a dynamic being reflected in time series data. Therefore, an ideal reconstruction algorithm should be able to estimate the functions *f*(*x*) and *g*(*x*), find the distribution of noise source ξ(*t*) (e.g., is it Gaussian or heavy-tailed?), assess the correlational properties of the noise (e.g., it is white or colored), account for the measurement errors, account for the issue of low-resolution in data, etc. We are not aware of a reconstruction algorithm which can tackle all such characteristics. Some reconstruction schemes are capable of tackling rather low-resolution time series data and can reveal the multiplicative nature of noise, though limited to Gaussian white noise (Aït‐Sahalia 2002b, Honisch and Friedrich 2011). Some can tackle the heavy-tailed character of noise distribution, though limited to white noise (Siegert et al. 1998, Aı and Yu 2006). And, some can cope with the colored nature of noise (Czechowski 2016, Lehle and Peinke 2018). In this paper we used two different reconstruction algorithms. The first one is designed to handle Langevin models in (8) under the assumption that the noise source ξ(*t*) is Gaussian and white but noise can be multiplicative (Siegert et al. 1998, Rinn et al. 2016). This algorithm, therefore, should be used whenever the dataset under study does not have big jumps. In practice, such an algorithm often works well for simulated data (see two simulated cases in Figure 5). In Figure 5A, you see two datasets with identical distributions and they look similar by eye. However, the underlying deterministic and stochastic mechanisms are completely different (see Appendix S5 for details). The reconstruction algorithm could successfully distinguish between the deterministic and stochastic components in the first dataset (Figure 5B&C) from those in the second dataset (Figure 5D&E). The second reconstruction algorithm we used is more advanced and can account for the presence of big jumps (Siegert and Friedrich 2001b) and is, therefore, more useful to reconstruct real datasets (see Appendix S3 for details). We first applied this algorithm to a simulated dataset for a proof of concept (Figure 3). We, then, applied the algorithm to an ice-core calcium record from the GRIP (Greenland Ice Core Project) (Fuhrer et al. 1993, Ditlevsen 1999b) (Figure 4).

## RESULTS

### When stochasticity is state dependent (Multiplicative noise)

A common assumption on stochastic processes in ecological models is that they result in random additions or removals of (bio)mass irrespective of the state. With such additive noise, the magnitude of stochastic fluctuations does not depend on the state of the system. For many real-world ecological systems this assumption is violated, as the magnitude of fluctuations depends on the state (called multiplicative noise. Figure 1B). For instance, due to demographic stochasticity, i.e., a type of stochastic fluctuation which occurs due to intrinsic mechanisms as a result of the discrete nature of birth and death processes in a population (Boettiger 2018), the relative magnitude of noise may be higher in smaller populations (DeAngelis and Mooij 2005). More precisely, in population dynamics models the magnitude of demographic stochasticity scales with the square root of the population size (Hakoyama and Iwasa 2000, Bonachela et al. 2012, Boettiger 2018). Environmental stochasticity, i.e., stochasticity which is driven due to external factors (Boettiger 2018), on the other hand, scales usually with the population size (Hakoyama and Iwasa 2000, Nolting and Abbott 2016, Boettiger 2018). But it is also possible that environmental fluctuations (e.g., climate) affect the parameters of the system (e.g., growth rate or mortality rate) resulting in a more complex effect on the state (Ripa and Heino 1999). For instance, fluctuations in any parameter of the grazing model of May (May 1977) generate stochastic regimes which are quadratic or more complex functions of the state (Appendix S2).

As a result, multiplicative noise can distort the stationary probability distributions. For instance, if the relative growth rate in the grazing model of May fluctuates by a white Gaussian noise, this translates to a quadratic multiplicative noise, which deforms the stationary distribution and displaces the modes. Since in this case the noise intensity near the under-grazed (high biomass) state is higher compared to the over-grazed (low biomass) state, the mode corresponding to the high biomass state shrinks with increasing noise and eventually disappears completely at a noise-induced bifurcation (Figure 2A). At even higher noise intensities the other mode can disappear as well. Constructed functions describing multiplicative noise can also deform the stationary probability distribution completely from unimodal to bimodal when the May model possesses only a single equilibrium (not shown).

**Figure 2.**
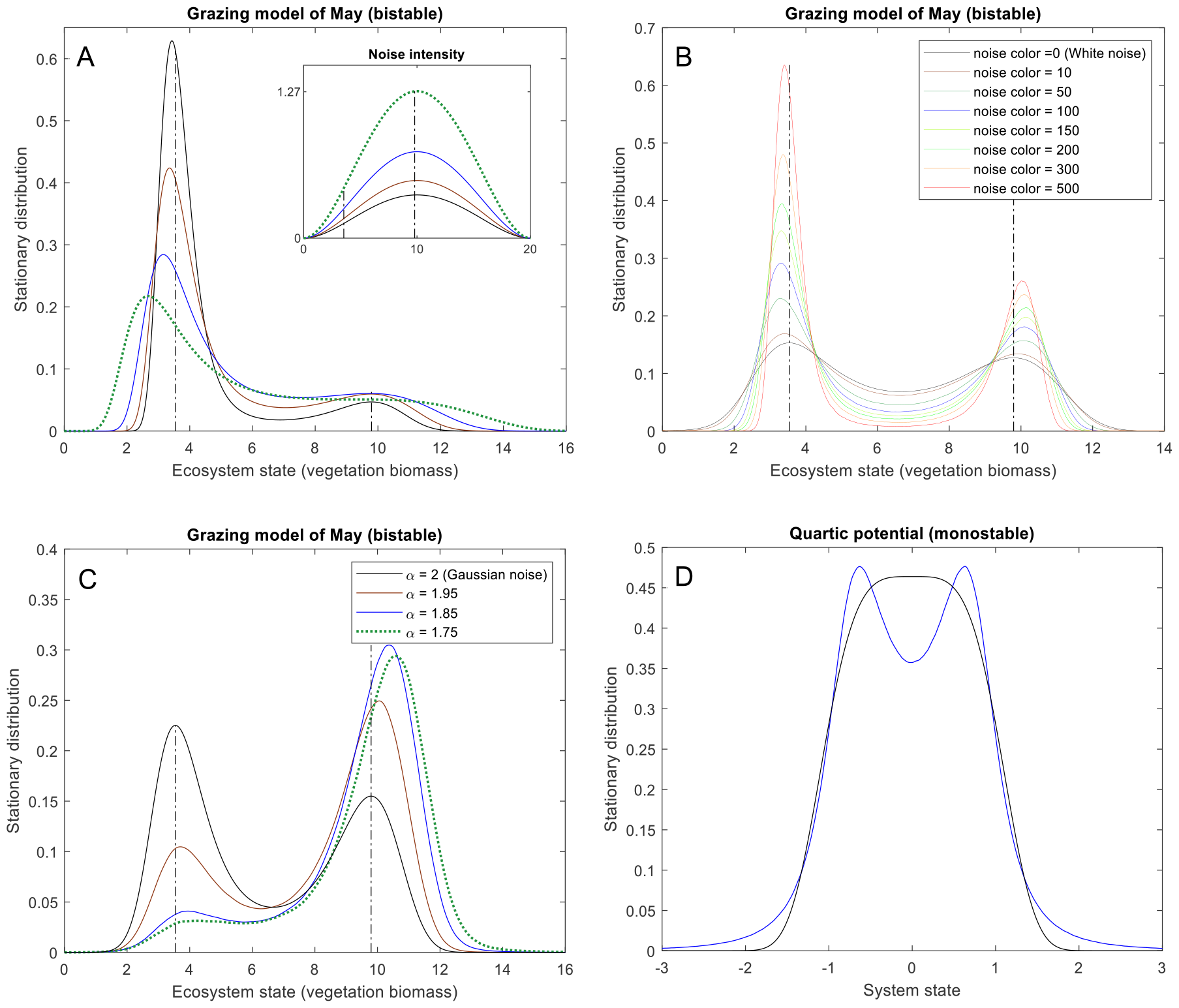
Effects of different kinds of noise on simulated frequency distributions. Modes can become displaced and hard to distinguish if noise is multiplicative (due to the variations of recovery rate in the May model (see Appendix S2)) (A), autocorrelated (B), or non-Gaussian (C). One of the two modes of the distribution can disappear (A and C) or a single-mode distribution can become bimodal (D). The dot-dashed lines represent the locations of the equilibria. For details of simulations see Appendix S6. See the beginning of appendix to reproduce this figure.

**Figure 3.**
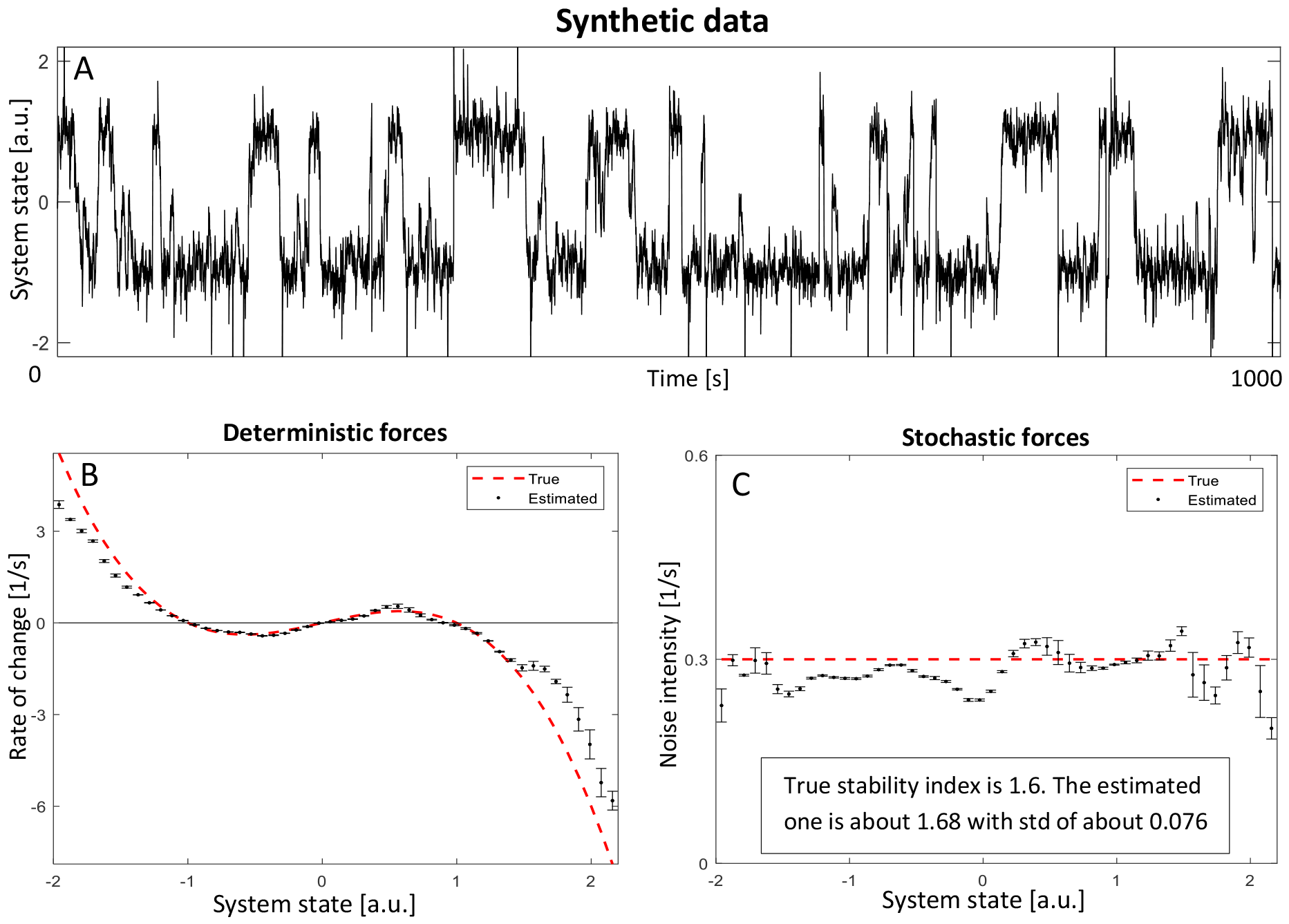
An example of how system reconstruction may infer the deterministic part and the noise from a time-series. We simulated a synthesis time series with 100000 data points from a simple bistable system being driven by an additive white Lévy noise. We applied the reconstruction algorithm in (29). The error bars are the corresponding uncertainties. For full details see Appendix S4. See the beginning of appendix to reproduce this figure.

**Figure 4.**
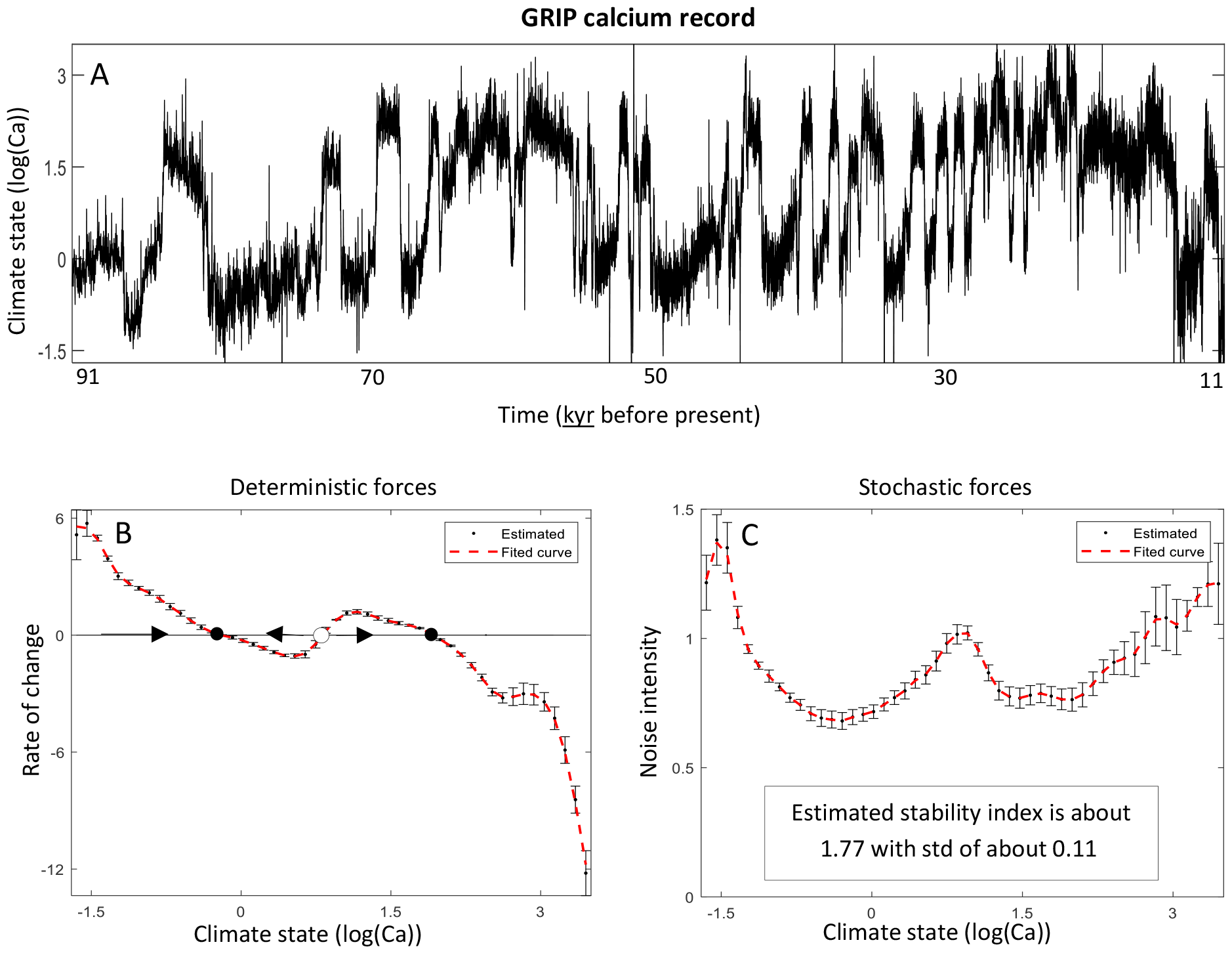
An example of how system reconstruction may infer the deterministic part and the noise from a time-series. We used calcium concentrations from the GRIP (Greenland Ice Core Project) record (data were originally published in (Ditlevsen 1999b)) (panel A) as a proxy for climate (Ditlevsen 1999a) during the last glaciation when the climate alternated between the cold glacials and warmer interstadials, a phenomenon called Dansgaard-Oescher events (Dansgaard et al. 1993a). Applying the reconstruction algorithm in (Siegert and Friedrich 2001a), the results indicate how the deterministic (black dots, panel B) and the stochastic (black dots, panel C) components of the dynamics varied with the state. The error bars are the corresponding uncertainties and the red dashed curves are smoothed functions going through after accounting for the uncertainties. The three zero-crossings rate-of-change curve in panel B suggests the existence of two alternative attractors (solid dots), and one repellor (open dot). The stability index was estimated to be 1.7877 suggesting a Lévy noise where extreme events are more common than expected from Gaussian noise (see the box in panel C and appendix S4). See the beginning of appendix to reproduce this figure.

**Figure 5.**
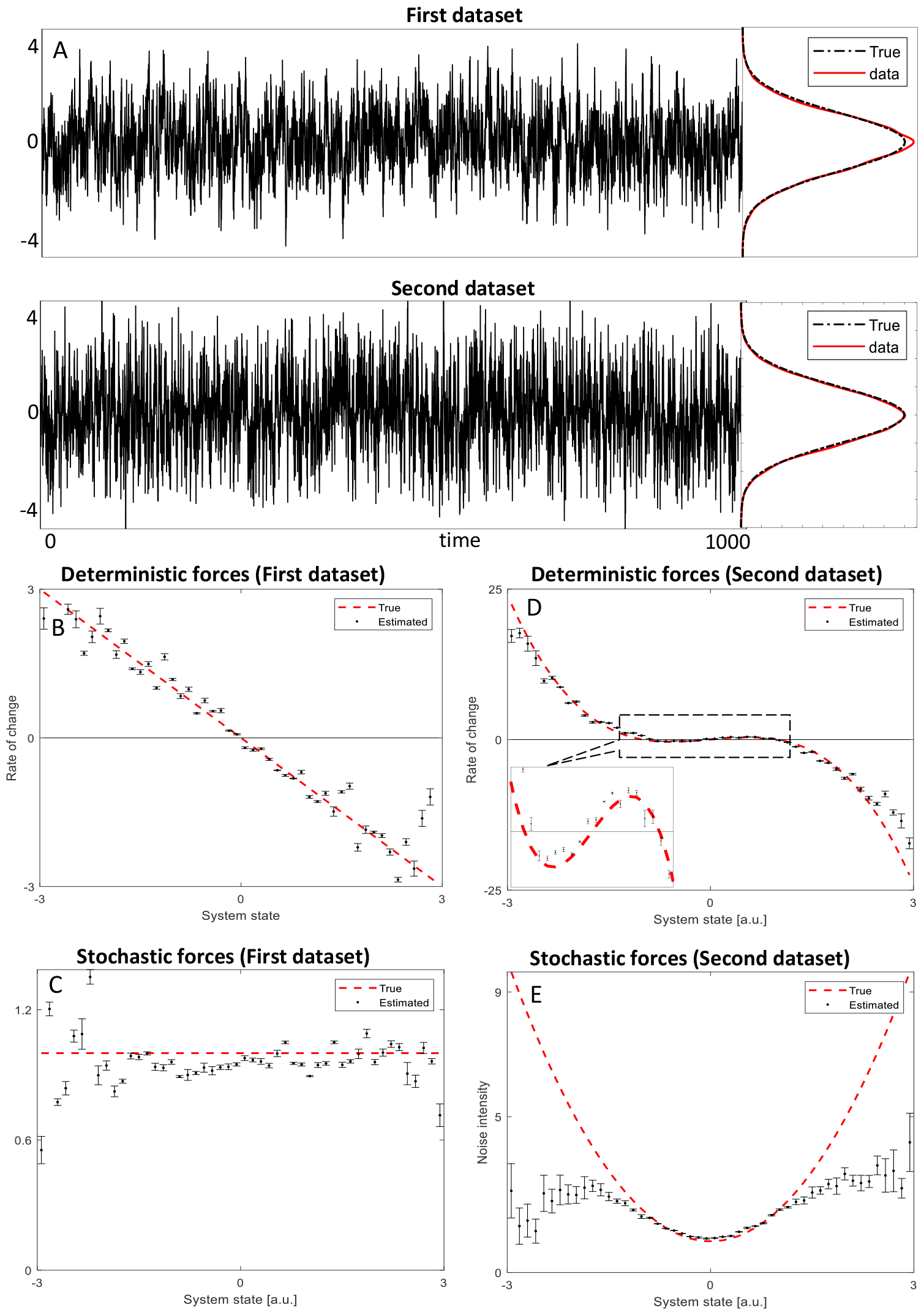
An example to clarify that disentangling different mechanisms in timeseries data is only possible through a proper system reconstruction. Two datasets with identical distributions (standard Gaussian distribution) being generated using different mechanisms (A). Reconstruction of the deterministic and stochastic parts for a dataset simulated from a process with a linear deterministic force (B) coupled to an additive noise (C). This should be compared with the reconstruction of another dataset simulated from a process with cubic deterministic laws (D) and a quadratic noise intensity (E). For details see Appendix S5. See the beginning of appendix to reproduce this figure.

### When extreme events predominate (non-Gaussian noise)

The Gaussian distribution is used often as the basic model describing stochastic perturbations. The logic behind this is simple: noise reflects our lack of knowledge about many unmeasured random forces of small impacts whose collective average effect, bearing in mind the Central Limit Theorem, can be described by a Gaussian distribution. The Gaussian distribution is, however, characterized by relatively short tails making extreme events unlikely (Figure 1C). On the other hand, many natural disturbances such as large storms, earthquakes, floods, and fires have a fat-tailed distribution (Carpenter et al. 2012) and resulting ecological forcing such as variations in annual nutrient loads to lakes can exhibit ‘jump-like’ behavior (Carpenter and Brock 2011, Carpenter et al. 2018). We can account for such rare events by using noise with a fat-tailed distribution based on a power-law (Figure 1D, also see Appendix S3). The effect of such jumps, the so-called ‘*Lévy flights*’, on the stationary probability distributions can be pronounced. This type of perturbation regime tends to shift the modes of stationary distributions, skew them, and shrink them as seen well by our analysis of the grazing model of May (Figure 2C). Further change of noise parameters (see Figure 2 and Appendix S3 for the stability index (parameter *α*)) can even make the probability distribution of the bistable model of May unimodal (dotted distribution in Figure 2C). On the other hand, Lévy noise can also evoke a bimodal state distribution in a system without underlying alternative stable states. This may be illustrated using a single-well quartic potential (a polynomial potential of degree four) whose stationary probability distribution is unimodal under additive Gaussian white noise (Dybiec et al. 2007). Stochastic fluctuations of Lévy flight type can change the unimodal probability distribution in this situation to bimodal (Figure 2D).

### When perturbations vary smoothly rather than sharply (Colored noise)

The forcing ‘noise’ in ecosystems typically comes from dynamical systems such as climate, the surrounding landscape or seascape, or unmeasured interactions that turn out to be important. Even if such dynamical systems themselves would be perturbed by white noise (uncorrelated), the resulting driving force they have on focal ecological systems will always be autocorrelated (Hanggi and Jung 1995) in the sense that the weather today is related to the weather of yesterday, and so on (Figure 1E and F). Such colored noise can also distort the stationary probability distributions. Physicists typically use the Ornstein-Uhlenbeck process as a noise source to study colored noise (Häunggi and Jung 1994). In this formulation the noise color is set by a parameter (*τ*, see Appendix S3) that raises the noise autocorrelation (i.e., it becomes redder) while at the same time decreasing the variance of noise. This combined change causes the system to stay nearer to its equilibria, making the probability distribution modes higher and narrower. Also, the position of the modes shifts (Häunggi and Jung 1994) in rather complex ways (Figure 2B, also see Appendix S3).

### Towards reconstructing the dynamical system and the noise simultaneously

Obviously, insight in the effect of noise characteristics is of limited practical value if we cannot determine the true character of stochasticity from data. Owing to the complex nature of unknown stochastic fluctuations this is not an easy problem even if one has complete knowledge about the underlying deterministic laws. However, there are ways to reconstruct the underlying deterministic laws and the character of the stochastic perturbation regime (the noise) simultaneously. Most of those reconstruction schemes require time series that are long enough to encompass multiple changes of state and sufficiently high-frequency to estimate variability. Even if such long and dense time series data are available there is no golden reconstruction scheme which takes into account all issues about the noise and can properly reveal the hidden structures and mechanisms. Reconstruction is a kind of inverse problem and in general such problems are not easy to tackle. Nonetheless, there has been exciting progress in this field during the last 25 years (Friedrich et al. 2000, Aït-Sahalia 2002, Aït‐Sahalia 2002a, Bandi and Nguyen 2003, Bandi and Phillips 2003, Friedrich et al. 2011, Honisch and Friedrich 2011, Tabar 2019).

The reconstruction algorithm in reference (Siegert and Friedrich 2001b), to the best of our knowledge, is one of the most advanced reconstruction algorithms and can reveal extreme perturbations typical in real datasets. To illustrate the usefulness and capacity of this algorithm we first applied it to a synthetic data simulated from a simple bistable system subject to additive white noise with a rather high jumping frequency (Figure 3A, see Appendix S4 for details). Although this pseudo dataset is moderately small the reconstruction technique detects deterministic and stochastic forces as well as the nature of stochastic jumps with a good accuracy (Figure 4 B&C). We also applied the reconstruction scheme to an ice-core calcium record from the GRIP (Greenland Ice Core Project) (Fuhrer et al. 1993, Ditlevsen 1999b) that has been analyzed before assuming Gaussian noise (Arani, et al. 2021). The results are consistent with the existing view that in the course of the last glaciation the climate shifted repeatedly between alternate climate regimes of cold glacial and warmer interstadial (Figure 4, Appendix S4), a phenomenon called Dansgaard-Oeschger events (Dansgaard et al. 1993b). However, this kind of analysis also suggests characteristics of the stochastic perturbation regime. This noise seems to have had a heavy-tailed distribution and forcing the climate in a multiplicative way (Figure 4C).

## DISCUSSION

Environmental sciences including ecology must draw inferences from data generated by deterministic and stochastic processes, including interactions of structure and stochasticity (such as multiplicative noise). Stochastic processes themselves may have complexities such as unbounded moments, autocorrelation, or complex modalities. We face this difficult problem with traditions that draw distinctions between environmental or external noise, endogenous noise, and noise co-created by our observational tools even though all sources of noise are present together. In this situation we are best served by datasets that are extensive in time and space with frequent observations and consistent methods comparable across years, locations, and different research groups, and analyzed by methods that account for deterministic and stochastic effects.

Here we have reviewed some of the challenges and existing solutions.

### Inferring alternative stable states from ecological data: caveats and challenges

As we have shown there is, in general, no one-to-one relationship between the stationary probability distribution and the underlying deterministic system. Due to different regimes of stochastic perturbations, systems with the same stationary distribution may have significantly different underlying deterministic laws (Figure 5). Therefore, analyzing the stationary distribution (e.g., frequency distribution analysis and potential analysis (Livina et al. 2010)) alone may lead to wrong conclusions about the existence and position of alternative stable states and ecosystem resilience (Scheffer et al. 2001). For instance, unimodal frequency distributions can arise even if the deterministic system has alternative stable states. It is also possible that complex noise can cause a deterministic system with a single stable state to produce a multimodal frequency distribution of states. If there is high-resolution data available, more sophisticated system reconstruction methods (Siegert et al. 1998, Tabar 2019) may help to reveal complex structures hidden in the data and to infer whether or not there are underlying alternative stable states.

An important limitation is that these system reconstruction methods need long time series data of high quality. Further investments may be needed to amass long-term, high-resolution ecological data. Long-term observation networks and paleoecology fill the critical need for long-term observations (Müller et al. 2010, Cowles et al. 2021). In some cases, extensive spatial data, e.g. from remote sensing, may allow ‘space for time’ substitution to infer alternate states from ecosystem change across gradients in conditions (Pickett 1989). Novel technologies are rapidly bringing accurate high-resolution collection of data within reach. For instance, high-frequency sensors in lakes and oceans and eddy flux techniques in terrestrial ecosystem allow for monitoring ecosystem processes at high temporal resolutions (Aubinet et al. 2012).

In conclusion, the ecosystem states and potential landscapes that we observe are the result of the interplay between deterministic forces and a stochastic regime of perturbations. Assuming the simplest regime of additive Gaussian white noise we may estimate the stability properties of the deterministic part directly from distributions of state. However, as we have shown, different kinds of stochastic regimes can confound the results. One could argue that separating noise from deterministic processes is artificial after all, and the distribution essentially reflects the true and effective dynamical properties of the whole. Nonetheless the system might settle into different stable states upon the removal of stochasticity (Figure 5). Novel approaches may pave the way to disentangle the role of noise and the underlying deterministic system. The prospects for such data-intensive methods may rise steeply as more high-density long-duration time series become available from satellite sensors and other novel technologies for automated observation.

In this paper, we showed that complex noise can impact the number, position and in general the distribution of the states partially or completely using one-dimensional stochastic systems. The outcome of such complex noise can lead to some particular phenomena such as extinction, persistence, coexistence and quasi-cycles in which the deterministic formalism cannot predict (Boettiger 2018). We also showed that such intricate mechanisms can only be correctly estimated upon the availability of high-resolution data by resorting to the state-of-the-art reconstruction algorithms. Here, we suggest several future research lines. One important future line of research is to investigate the impact of complex noise in more complex high-dimensional systems and see how the curse of dimensionality comes into play. Likewise, reconstruction algorithms need to be developed to be able to tackle multivariate time series of observations as the current algorithms might perform poorly in high dimensions. One particularly interesting and elegant class of complex high-dimensional systems is network dynamical systems which can model important systems like pollinator-plant systems. To the best of our knowledge reconstruction algorithms are so far limited to reconstruct the deterministic rules of networks. Furthermore, we mentioned in this paper that high resolution data are needed to estimate the underlying stochastic system reliably. However, there are situations in which high-resolution data are not available. Therefore, parallel to putting effort to get high-resolution measurements it is of great importance to also develop reconstruction algorithms which can fill the gaps for the sparsely sampled observations (currently, there are some algorithms (Aït‐Sahalia 2002a) to cope with low-resolution data but they have their own limitations). Finally, it is important to develop algorithms which can tackle extensive spatial and temporal observations.

## Acknowledgments

S.R.C. is supported by U.S. NSF grants DEB-2318567 and DEB-2025982. C.X. is supported by the National Natural Science Foundation of China (31770512) and the Fundamental Research Funds for the Central Universities (020814380089).

## Supplementary Information

**Note:** You can find the MATLAB codes and reproduce Figures 2-5 in Github repository https://github.com/mshoja/Reconst (Babak M. S. Arani 2023) by running the main code called AllFigures.m.

## Appendix S1 Correspondence between multimodality and multistability under additive Gaussian white noise

In this appendix we show that in a stochastic model with one state variable and additive Gaussian white noise, there is a one-to-one correspondence between the local maxima (i.e., the modes) of the stationary probability distribution and the stable equilibria of the deterministic component of the system (Likewise, the minima of the stationary distribution correspond to the unstable equilibria). In higher dimensions our proof holds for systems that fulfil the so-called ‘potential condition’ (Risken 1996) with a diagonal diffusion matrix where the system resembles a one-dimensional system with a similar proof. We try to give a very simple proof.

Consider a stochastic process *x*(*t*). Its change in time, 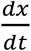, can be expressed by both a deterministic force *f*(*x*) and stochastic fluctuations ξ(*t*) (i.e., noise) via the so-called ‘Langevin equation’

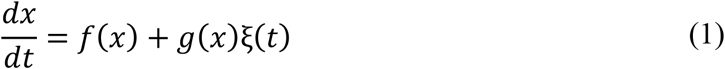

Where the function *g*(*x*) actually weighs the stochastic fluctuations ξ(*t*) (note that in this paper we adopt an Ito interpretation for the stochastic integration). If *g*(*x*) is independent of the state of the system (i.e., *g* is constant), the noise is called additive. Otherwise, it is called multiplicative. The stationary probability distribution of (1) (*p*^*st*^(*x*)) under Gaussian and white noise ξ(*t*) (ξ is standard Gaussian) is (Risken and Risken 1996)

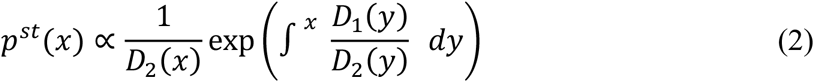

Where *D*_1_(*x*) = *f*(*x*) and 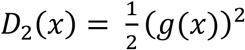 are called ‘drift’ and ‘diffusion’ coefficients and *∝* denotes proportionality. Since we have an additive noise *D*_2_(*x*) = *D*_2_ = const, the relation (2) simplifies to 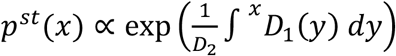. We note that the integral − ∫^*x*^ *D*_1_(*y*)*dy* equals the potential function *U*(*x*) = −∫^*x*^ *f*(*y*)*dy*, so we can write:

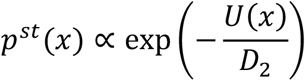

Since the exponential function exp(x) is an increasing function, we know that the minima and maxima of *p*^*st*^(*x*) and 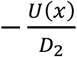 are the same. Due to the negative sign in 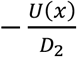 (note that *D*_2_> 0) the maxima and minima of *p*^*st*^(*x*) correspond to the minima and maxima of the potential *U*(*x*), respectively. It should be clear to see that the minima of our potential (downhills) actually correspond to the stable equilibria and the maxima of the potential (uphills) correspond to the unstable equilibria. This completes the proof.

## Appendix S2. Additive noise on parameters usually leads to multiplicative noise in non-linear systems

In this appendix, we show that additive noise on a parameter of a deterministic system is usually translated to a stochastic system with a multiplicative noise (Ripa and Heino 1999). As an example, we study the additive perturbations on the parameters of the grazing model of May (May 1977) (see Appendix S6 for details about the May parameters). Assume that the relative growth rate (r) fluctuates by an additive white noise term ξ(*t*), i.e., *r* ← *r* + ξ(*t*). Inserting this perturbed growth term in the original deterministic model of May clearly leads to the following stochastic system with a multiplicative noise of quadratic type

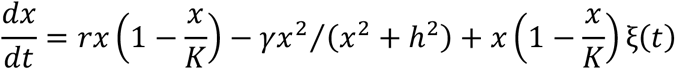

If, however, the carrying capacity (K) fluctuates, *K* ← *K* + ξ(*t*), we cannot factor out the original deterministic part. In this case, one can find the following Langevin approximation of the resulting stochastic system via a Taylor series expansion of 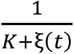 about the mean value of perturbations (which is 0) by keeping only the first two terms

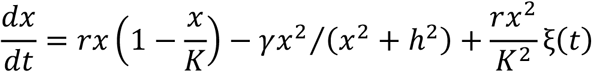

In general, assuming *f*(*x, λ*) is differentiable with respect to the second argument, one can find the following Langevin approximation (up to the linear term) to the general model 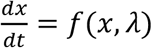 when the parameter *λ* fluctuates (*λ* ← *λ* + ξ(*t*))

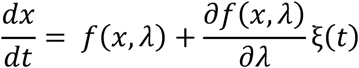

Where the noise is clearly multiplicative if 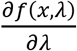 depends on the state variable *x*. Phrased differently (and assuming *f*(*x, λ*) is differentiable with respect to both arguments), the noise is additive if and only if the function *f*(*x, λ*) can be separable in terms of two functions purely in terms of *x* and *λ*, i.e., *f*(*x, λ*) = *f*_1_(*x*) + *f*_2_(*λ*).

## Appendix S3. A short description about Lévy (or alpha-stable) noise and coloured noise

In this appendix, we describe the different kinds of noise used in the main text in some details and explain how they are used in our simulations.

### White Gaussian noise

Thinking of noise as cumulative effect of many variables (degrees of freedom) the Gaussian distribution is a typical candidate for the distribution of noise due to the Central Limit Theorem (CLT). CLT asserts that the sum of a sequence of independent and identically distributed (iid) random variables with ‘*finite variance*’ converges to a Gaussian distribution even if the original variables themselves are not Gaussian distributed. The white aspect of noise (lack of temporal correlations) reflects a ‘*fast*’ fluctuating environment, i.e., the contributing degrees of freedom operate at short time scales, and relates to a timescale separation between the system and the noise.

### White Lévy noise

In real world problems the assumption of finite variance for the contributing variables to the noise can be violated due to the presence of extreme events in which we need to resort to the generalized CLT. In such cases the sum of iid variables will tend, instead of Gaussian, to an *α* − stable distribution with *α* < 2. Therefore, *α* − stable distribution is a more realistic candidate to account for the distribution of noise. This distribution is a generalization to the Gaussian distribution and has four parameters: 0 < *α* ≤ 2 (stability index), -1 ≤ *β* ≤ 1 (skewness parameter), *μ* (location parameter), and *σ* (scale parameter). Note that the Gaussian distribution is an extreme member of the *α* −stable family of distributions and corresponds to *α* = 2. For other ranges of the stability index (0 < *α* < 2), the *α* −stable distribution is fundamentally different from the Gaussian distribution: unlike the Gaussian distribution where tails decay very fast (exponentially decaying tails), the *α* −stable distribution with *α* < 2 has ‘*heavy*’ tails which asymptotically follow the power law. More precisely, the tail(s) tend to |*x*|^−(1+*α*)^ for large *x*. The parameter *β* controls the asymmetry (skewness) of the distribution so that -1 ≤ *β* < 0 corresponds to negative skewness, 0 < *β* ≤ 1 corresponds to positive skewness, and *β* = 0 corresponds to a symmetric *α* −stable distribution. We chose *μ* = 0 since a non-zero location parameter can be expressed in the deterministic part of the system. The scale parameter *σ* controls the dispersion of the distribution (although the variance of a stable distribution is undefined for *α* < 2).

### Colored Gaussian noise

Unlike white noise where the autocorrelation function decays immediately to zero, the autocorrelation function of colored noise decays gradually. This implies that in systems driven by colored noise information about the history is needed to predict future states. This, therefore, makes such systems difficult to study. For simplicity, it is common to use the Ornstein-Uhlenbeck process as colored noise source (therefore although the system has memory of the past the noise itself has a minimal memory, i.e., noise is a Markov process). So, to study systems driven by colored noise we assume that the noise source in (1) evolves as

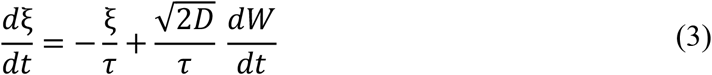

where *W* stands for the standard Wiener process (Brownian motion) and the correlation function of noise decays exponentially

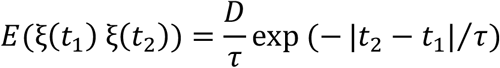

Unlike white noise which has only one parameter (i.e., noise intensity *D*) colored noise has a second parameter (*τ*) called ‘*noise correlation’*.

It is interesting to note that in our simulation of grazing model of May (similar to the case in (Hanggi and Jung 1995), see Figure 6.4) driven by a colored noise the modes of stationary distribution shift as noise correlation increases reaching the maximum shift followed by a decrease for further increase of noise color (see Figure 3 in the main text). As noise color tends to infinity, the stationary density modes shift back to the equilibria again (similar to the white noise case) but now the peaks are extremely narrow.

### Estimating stationary distributions

In contrast to systems driven by Gaussian white noise, systems driven by Lévy noise or colored noise have a much more complex equation (called master equation) to describe the evolution of the probability distribution function. In the case of Lévy noise, this equation is a fractional Fokker-Planck equation whose numerical solution is slow and not stable (Dybiec et al. 2007). For systems driven by colored noise there is even no closed form master equation in general (Häunggi and Jung 1994). Therefore, we used Monte-Carlo simulations to estimate the stationary distributions when the noise is coloured or Lévy. Full details of simulations can be found in appendix S6.

## Appendix S4. Application of ‘system reconstruction’ using an ice-core climate record during the last glaciation

If we know all governing laws of a dynamical system, we can easily generate data by simulation. However, uncovering unknown governing laws through data only, is notoriously difficult. System reconstruction methods are recipes for this so-called inverse problem where both the deterministic and stochastic rules are found based on a time series. Here, we explain the method that we used in more detail.

### The data set

We analyzed the calcium record from the GRIP ice-core (Fuhrer et al. 1993). This time series has the highest temporal resolution (almost annual spanning from 11000 to 91000 years before the present) among glacial climate records (Fuhrer et al. 1993). The logarithm of calcium record (Figure 4A in the main text) serves as a climate proxy, which is highly stationary with a white but non-Gaussian noise source (Ditlevsen 1999c). Data in Figure 4A in the main text were originally published in as Figure 1 based on core depth versus concentration data presented by Fischer et al (2022) (see the link https://doi.org/10.1594/PANGAEA.942777) and the ice-core chronology in (Rasmussen et al. 2023). Professor Peter Ditlevsen kindly provided the data from his figure in tabular form.

### Pre-investigations on data

Prior to applying system reconstruction some pre-investigations on data are necessary to see if data fulfil the conditions needed by the reconstruction procedure we want to follow. However, it might still be possible to apply the reconstruction procedure if some such conditions are violated (Rinn et al. 2016b).

First, data should be stationary, meaning that the statistical properties of the data should remain unchanged in the studied period. Normally, a weak sense of stationarity is checked, i.e., one checks if the mean and the variance of time series remain unchanged and the autocorrelation function depends only on the time lag in the studied period. The reconstruction can, however, be applied to non-stationary data by a moving window technique in which the system is assumed to be quasistationary within each window. Obviously, this is only possible if we have enough data.

Second, most reconstruction methods assume a white noise source which implies that the future state of the system depends only on the present state (called Markov property). If this assumption is violated then the system has memory of its past and therefore, we need to use a sparser subset of data with a coarser time resolution under which memories of the past are removed (the so-called Markov-Einstein time scale) (Friedrich et al. 2011, Tabar 2019). This, again, might leave us with insufficient data. Our data set fulfills the mentioned criteria (Ditlevsen 1999b).

### The reconstruction scheme we used

Here, we outline the reconstruction algorithm in (Siegert and Friedrich 2001a, Siegert and Friedrich 2001b) and we constrain the discussion to univariate systems (we keep the same notations). This scheme requires the noise to be white and can reveal the multiplicative nature of noise and also can account for the presence of extreme events by replacing the Gaussian noise ξ in (1) by an *α*-stable noise ξ_*α*_

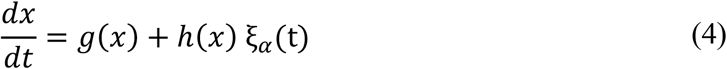

where ξ_*α*_ is a symmetric *α*-stable noise with zero mean and a scale parameter of 1 (μ = 0, σ = 1, β = 0) and 1 < *α* ≤ 2. Note that the noise term ξ_*α*_ reduces to Gaussian if *α* = 2. So, the *α*-stable noise ξ_*α*_ is a generalization to Gaussian noise ξ in (1) and based on the stability index *α* can have a heavy tail distribution (the smaller the *α* is the heavier the tails would be). Based on univariate time series data the following functions and parameters are estimated: the deterministic part *g*(*x*), the stochastic part *h*(*x*), and the noise parameter *α*.

*g*(*x*) and *h*(*x*) are unknown (most probably) non-linear functions of the state *x*. To describe the shape of these functions we discretize the values of the state variable *x* into bins. Here, a balance between the number of data points and bin size is important to make sure that there is enough data per each bin and that there is enough number of bins to adequately describe the functions (we considered 50 bins).

The procedure first tackles the estimation of the deterministic part *g*(*x*) for each bin *x* (we use the same notation *x* to refer to bin centres. So, by bin *x* we mean the bin whose centre equals *x*)

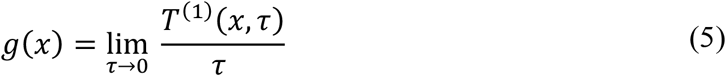

where the numerator is the conditional average *T*^(1)^(*x, τ*) = *E*(*x*(*t* + *τ*) − *x*(*t*))│_*x*(*t*)=*x*_. The meaning of the condition *x*(*t*) = *x* in *T*^(1)^(*x, τ*) for our ‘*discrete*’ time series data *x*(*t*_*n*_), n=1, 2, … is that only the differences *x*(*t*_*n*_ + *τ*) − *x*(*t*_*n*_) in which *x*(*t*_*n*_) is within the bin *x* are considered. This condition is expressed as *x*(*t*_*n*_) = *x* ± Δ*x* in bellow where Δ*x* is half bin size.

Then *T*^(1)^(*x, τ*), for a fixed bin *x*, can be estimated as

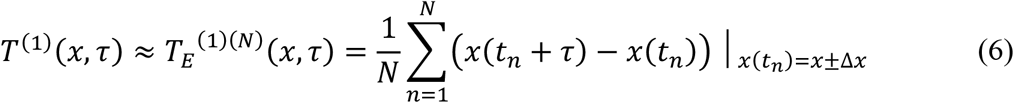

where *N* is the amount of data in bin *x*. In *T*_*E*_^(1)(*N*)^(*x, τ*) the subscript *E* is added to emphasize that it is an estimation to *T*^(1)^(*x, τ*) and the superscript *N* is added to stress the dependency of *T*_*E*_ on the amount of data in bin *x* (a large *N* is needed for a good estimation).

Furthermore, in (5) the limit of *τ* → 0 is needed. So, in practice we should calculate *T*_*E*_^(1)(*N*)^(*x, τ*) for ‘*small*’ values of *τ*, i.e., for some few first multiples of the sampling time Δ*t*_sample_ (*τ* = Δ*t*_sample_, 2Δ*t*_sample_, 3Δ*t*_sample_, …) and such values of *τ* should be much smaller than the (unknown) relaxation time of the system *τ*_*R*_. One can roughly estimate *τ*_*R*_ by fitting the autocorrelation function of the Ornstein-Uhlenbeck process (viewed as a linear approximation to our unknown nonlinear system), i.e., the exponential *e*^−*t*/*τ*^*R*, to the autocorrelation function of data. Doing so, we find that *τ*_*R*_ ≈ 1000 years. We, then, considered only the first five sampling times (*τ* = Δ*t*_sample_, …, 5Δ*t*_sample_) and since the sampling time is annual our choice makes sense.

Now, for a fixed bin *x* the deterministic part of the system, i.e., *g*(*x*), can be estimated by extrapolation of *T*_*E*_^(1)(*N*)^(*x, τ*) values (*τ* = Δ*t*_sample_, …, 5Δ*t*_sample_) to *τ* = 0 as the limit *τ* → 0 is needed in (5). This gives us an estimation 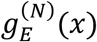 for the deterministic part. Following the ideas in (Rinn et al. 2016a) we estimated *g*(*x*) from the slope of a ‘*weighted*’ linear regression line to *T*_*E*_^(1)(*N*)^(*x, τ*) values for *τ* = Δ*t*_sample_, …, 5 Δ*t*_sample_ (we describe how to find the weights, i.e., the error bars, later). The algorithm we are explaining is an ‘*iterative*’ algorithm: in the first iteration we do not know the weights so in this first step unit standard deviations is used as weights (unweighted regression). Once an estimation of the system is at hand we can estimate the uncertainties of the results, i.e., the weights. Afterwards, we can update the results using the weights, then update the weights again using the new results, update the new results using the new weights, so on. Actually, the first iteration gives us an already good result and in practice the algorithm should almost converge after two iterations (we considered four iterations).

The next step is to estimate the stability index, i.e., the parameter *α*. The following relation

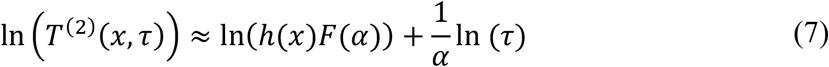

gives us the clue where *T*^(2)^(*x, τ*) is defined as

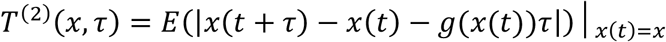

Which can be estimated in a manner similar to *T*^(1)^(*x, τ*) as

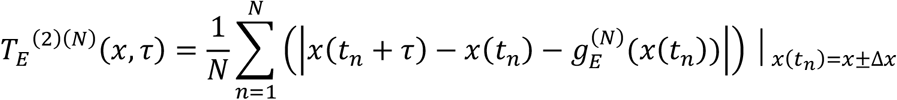

and *F*(*α*) can be estimated by simulating a long Lévy motion with *α* as the stability index and then taking the average of the absolute value of the realization. The simulation of a Lévy motion is described in (Siegert and Friedrich 2001a, Siegert and Friedrich 2001b). In MATLAB it can be done simply by the following commands

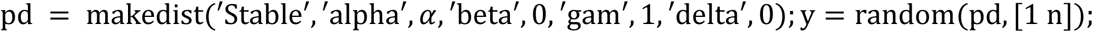

where *n* is the length of the realization *y* we like to get. By relation (7), one finds that for a fixed bin *x, α* can be estimated as the inverse slope of a fitted line to ln (*T*^(2)^(*x, τ*)) values versus ln (*τ*) values (Note that here we do not need to know the expression ln(*h*(*x*)*F*(*α*)) in (7) as this is the intercept of the mentioned fitted line. However, later we need to know *F*(*α*) for the rest of the calculations).

In theory we should get the same value of *α* for each bin *x* but in practice we get different values mainly due to finite data we have and also different amount of data in different bins, etc., This suggests to take the average of all values of *α* as an estimation of the stability index (*α*_*E*_) and their standard deviation as the uncertainty of the stability index Δ*α*_*E*_.

The last step concerns the estimation of the stochastic part *h*(*x*)

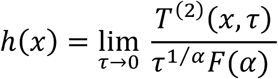

which can be estimated as

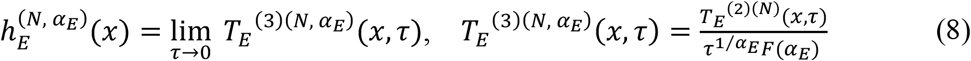

where the superscript *α*_*E*_ is added to remind that the estimation in (8) is influenced by the estimated value *α*_*E*_. The details of the calculations are exactly similar to the deterministic part.

Now, we describe calculations needed for the uncertainty analysis (calculation of error bars). In (6), the uncertainty of *T*^(1)^(*x, τ*) can be estimated as

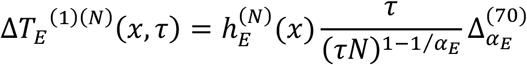

where the width 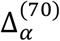 is defined in the following to deal with the uncertainty of a Lévy motion *Z* with *α* as the stability index (simulated in MATLAB as described above)

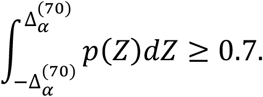

The rationale behind such a definition stems from the fact that for a Lévy motion the variance is not defined and we cannot proceed with the same way as was done with Δ*α*_*E*_. In (7), the uncertainty of ln (*T*^(2)^(*x, τ*)) is estimated as

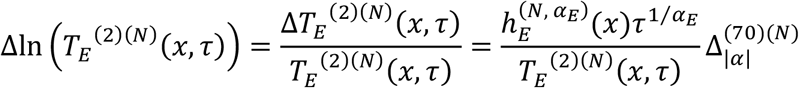

where the width 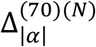 is defined in the following to deal with the uncertainty of the average of the absolute value of Lévy motion realizations *Z*, i.e., 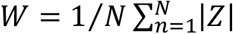

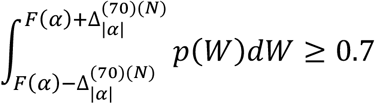

In (8), the uncertainty of *T*_*E*_^(3)(*N*)^(*x, τ*) can be calculated as

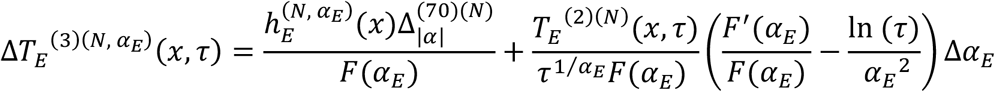

Finally, the statistical uncertainty of the deterministic part for a fixed bin *x*, i.e., *σ*^2^_*g*_(*x*), (see the error bars in Figures 3B and 4B in the main text) can be calculated as the uncertainty of the slope of the weighted regression line to the *T*_*E*_^(1)(*N*)^(*x, τ*) values for *τ* = Δ*t*_sample_, …, 5Δ*t*_sample_ (with Δ*T*_*E*_^(1)(*N*)^(*x, τ*), *τ* = Δ*t*_sample_, …, 5 Δ*t*_sample_, as the weights). The error bars of the stochastic part (see the error bars in Figures 3C and 4C in the main text) are calculated similarly and the uncertainty of the estimated stability index, i.e., Δ*α*_*E*_, is taken to be the standard deviation of all stability index estimated for all bins (see the boxes in the bottom of Figures 3C and 4C in the main text). For a few bins close to the edges of the data range, where there is much less data in the bins, we noticed that the stability indices were bigger than 2. We have excluded such values in the estimation of stability index and its uncertainty. Our calculations show that the stability index is estimated to be *α*_*E*_ ≈ 1.7877, very close to the value 1.75 in (Ditlevsen 1999c), with the uncertainty of Δ*α*_*E*_ ≈ 0.1037.

## Appendix S5. Different stochastic systems can have the same stationary distribution

Here, we consider two different systems: one being monostable driven by an additive Gaussian white noise while the other being bistable but driven by a multiplicative noise. For the monostable system we consider the Ornstein-Uhlenbeck process, i.e., Eq. (3) and set *D* = *τ* = 1 (with a realization illustrated in Figure 5 in the main text, see the time series in bottom left). The corresponding drift and diffusion functions are *D*_1_(*x*) = −*x, D*_2_(*x*) = 1, respectively. Using (2) the stationary distribution is proportional to

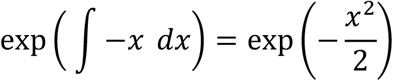

i.e., the stationary distribution is the standard normal distribution. On the other hand, for a typical bistable system with drift *D*_1_(*x*) = *x* − *x*^3^ driven by a multiplicative noise of quadratic type with *D*_2_(*x*) = 1 + *x*^2^ (with a realization illustrated in Figure 5 in the main text. See the time series in bottom right) the stationary distribution is proportional to

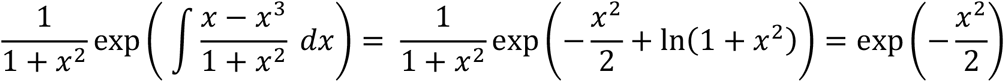

i.e., the stationary distribution is again the standard normal distribution. This means that the stationary effective potential for both systems is the same but they clearly have different natures.

## Appendix S6. Simulations in Figure 2

We used the grazing model of May in all figures (except in Figure 2D). The dynamics of the May model are defined by the following differential equation:

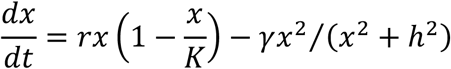

Where the state variable *x* represents the vegetation biomass; parameters *r, K, γ*, and *h* represent the relative growth rate, carrying capacity, herbivore’s consumption rate, and herbivore’s half saturation constant, respectively. Our choice of model parameters is *r* = 0.1, *K* = 20, *h* = 3.4, and *γ* = 0.56. For this parameter set the May model is bistable. In Figure 2D, the model is designed to have a potential, *U*(*x*), which is a polynomial of degree four (quartic potential), hence the dynamics follows a cubic polynomial 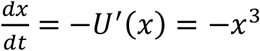.

All stationary probability distributions in which the underlying noise is white and Gaussian (additive or multiplicative) are calculated by numerically solving the corresponding Fokker-Planck equations (we used the partial differential equation solver *pdepe* in Matlab 2011b). A Monte-Carlo simulation (Euler-Maruyama scheme) is used to estimate the stationary probability distributions when noise is colored or Lévy. More specifically, we simulated many long realizations in a parallel manner (see Table S6) instead of one extremely long realization to speed up the calculations. The choice of the number and length of the realizations are made by trial and error based on the fact that stationarity should be reached. After that, we discarded the first 10% of all trajectories to make sure that dependence on the initial conditions is gone. Then, we used every 100 points in each trajectory and afterwards used the Matlab code *kde*.*m (Botev et al. 2010a, b)* to estimate the stationary distributions. Following (Ditlevsen 1999c, a), for the case of Lévy noise we used Heun’s integration scheme since large excursions made by Lévy noise requires a more stable integration procedure for the drift term. We used integration time step Δ*t* = 10^−2^ in all simulations except for the case of quartic potential where a rather small-time step is needed for numerical accuracy (we used Δ*t* = 10^−3^, see (Dybiec et al. 2007)). There are some difficulties regarding the simulation of the May model under stochastic perturbations. Clearly, this model can make excursions to negative values of biomass due to noise. To avoid such a biologically impossible situation, we chose parameter settings under which the smallest equilibrium (i.e., 3.54) is rather far from zero. However, the system still can sometimes make excursions to negative values of biomass, especially under Lévy noise. Therefore, we used a reflecting boundary at 0 so that once the system hits 0 it will be reflected back into positive states. To address this, we used a reflecting boundary condition at zero in the implementation of the corresponding Fokker-Planck equations. In simulations, we used a rather simple *projection* method (it simply projects the trajectories back into domain boundary once they leave the domain). We also used a right reflecting boundary at 20 which is far enough from the greatest equilibrium (i.e., 9.79) in the implementation of Fokker-Planck equation. In the case of Lévy noise extra care is necessary. First of all a potential with edges steeper than a quadratic potential is needed to get bounded solutions for Lévy stability index *α* < 2 (Dybiec et al. 2007). This is why we used a quartic potential. Fortunately, the right edge of the potential for the May model is (asymptotically) steeper than the quadratic potential. Second, even if the potential is steep enough the noise can still make rather large and unrealistic excursions. Such very large excursions can lead to numerical inaccuracies no matter how small is our integration time step Δ*t*. To fix such a problem one should consider a cut-off so that the system is not allowed to go beyond it (called *truncated Lévy flights* (Méndez et al. 2013, 2016)). Our cut-off for the quartic potential was ±40. For the May model 0 was clearly our left cut-off and we chose 20 to be the right cut-off. Finally, for the sake of comparability between Gaussian noise and Lévy noise in Figure 1D, the scale parameter of Lévy noise is chosen to be 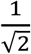 multiplied by the corresponding white noise intensity. The reason is that the standard deviation of Lévy noise with scale parameter *σ* (and *α* = 2) is 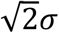. The following table summarizes the full details about the noise properties in Figure 2 (main text).

**Table S6.** Details on the noise properties in Figure 2 (main text).

|  | Noise characteristics | Noise parameters | Method of calculation |
| --- | --- | --- | --- |
| Fig. 2A | multiplicative Gaussian white | The only noise parameter is noise intensity which is 0.03, 0.04, 0.06, and 0.102 for black, brown, blue, and green distributions, respectively. | All distributions are calculated by solving the corresponding Fokker-Planck equation numerically. |
| Fig. 2B | additive Gaussian coloured | Noise has two parameters. Noise intensity is $\sigma=0.02$ for all distributions while noise autocorrelation varies (see the legends). | 1500 trajectories of length $10^6$ are simulated for the noise colors $\tau = 500$, $\tau = 300$, $\tau = 200$, $\tau = 150$, $\tau = 100$, $\tau = 50$, and $\tau = 10$ respectively. |
| Fig. 2C | additive $\alpha$ –stable white | Only stability index ($\alpha$) changes (see the legend) while other parameters of noise are kept fixed (skewness parameter $\beta = 0.03$, location parameter $\mu = 0$, and scale parameter $\sigma = 0.01$). | For $\alpha = 2$ (Gaussian noise) the corresponding Fokker-Planck equation is solved numerically. For $\alpha = 1.99$, $\alpha = 1.95$, and $\alpha = 1.70$ we simulated $10^3$, 800, and 600 trajectories of length $5 \times 10^6$. |
| Fig. 2D | additive $\alpha$ –stable white | In black distribution the only parameter is noise intensity which is 0.5. In blue distribution stability index $\alpha = 1$, skewness parameter $\beta = 0$, location parameter $\mu = 0$, and scale parameter $\sigma = \frac{1}{\sqrt{2}}0.5$. | For the black distribution the corresponding Fokker-Planck equation is solved numerically. For the blue distribution $10^5$ trajectories of length $2 \times 10^4$ are simulated. |

